# A unique two-excipient heat stable tablet formulation of oral insulin: preliminary results

**DOI:** 10.1101/2025.01.29.635377

**Authors:** Manshun Lai, Changcheng Zhu, Jaclyn Delarosa, Manjari Lal

## Abstract

We report a novel formulation approach for development of a thermostable oral insulin tablet. Using freeze drying to form a heat stable tablet in a single-step process, we demonstrate hydroxypropyl beta cyclodextrin (HP-β-CD) encapsulated lipophilic ion-pair complex of insulin using bile salt achieves intestinal absorption and sustained glucose levels. The tablets produced using this simple approach with only two excipients offer protection from enzymatic and stomach acid degradation and facilitate insulin uptake, without any need for specialized drug manufacturing or enteric coating. Insulin in this innovative formulation is thermotolerant, capable of maintaining stability even under heat stress at 30-40°C/65-75% RH. The convenient presentation of insulin as a thermostable oral tablet presents a low-cost scalable manufacturing method that simplifies the logistics of storage, transport, and distribution in any setting, including areas where cold storage maybe limited or unavailable.

**Graphical Abstract:** Our unique formulation approach protects insulin from stomach and temperature induced degradation, improves intestinal permeability providing a sustained release of insulin for glycemic control. The freeze dried tablet product format is easily scalable and flexible, increasing accessibility across all populations.

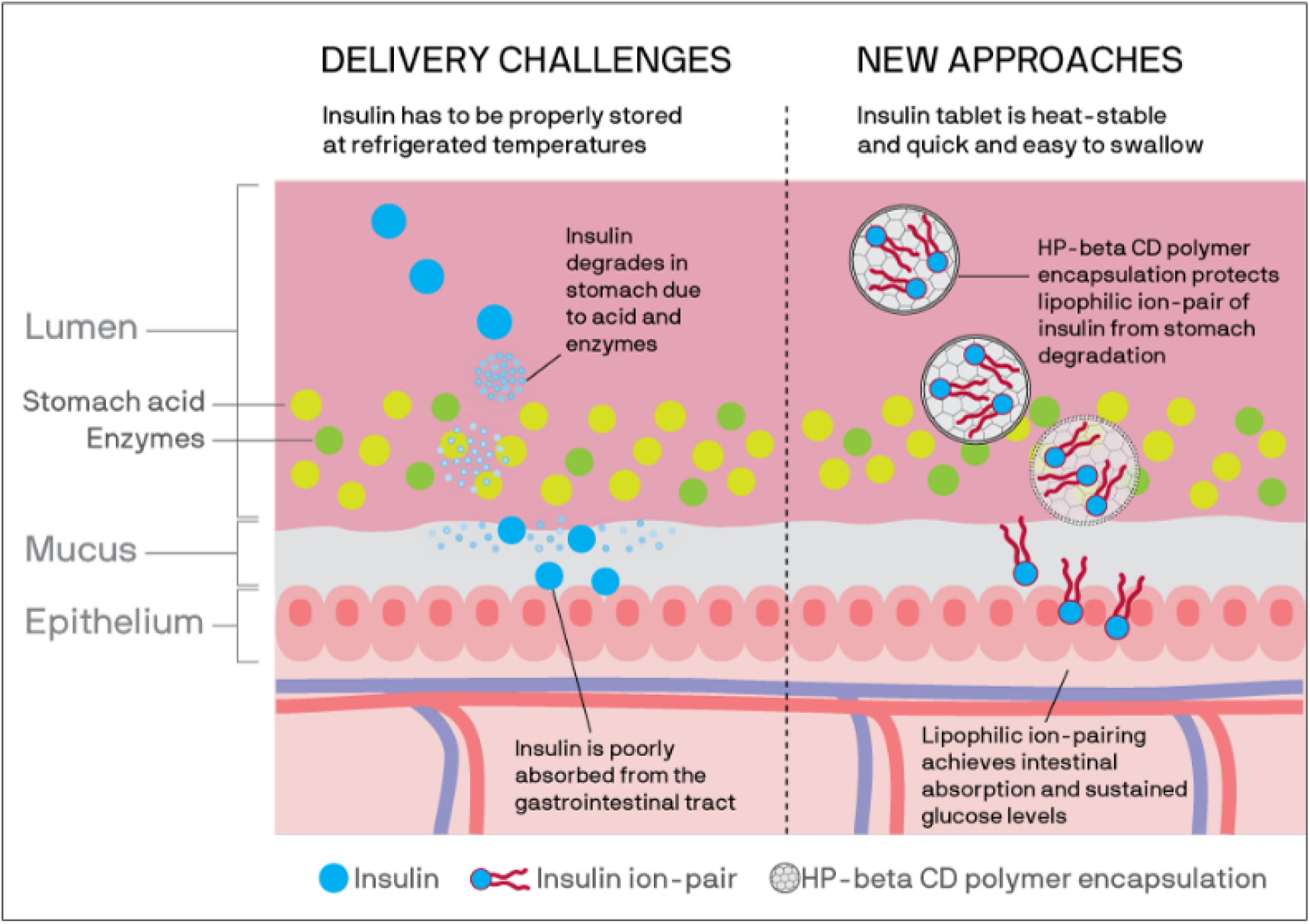

## Introduction

Globally, insulin remains underused due to barriers that hinder adherence, such as the well-known difficulties of frequent injections [1]. Degradation of insulin at both elevated and frozen temperatures makes it difficult to distribute and store in areas with limited or no access to refrigeration and a temperature-controlled supply chain. Oral insulin is attractive as it represents the potential for improved accessibility, acceptability, and dose adherence; however, in addition to gastrointestinal (GI) degradation, the high molecular weight and hydrophilicity of insulin limits intestinal absorption.

Previous oral insulin approaches using nanoparticles, nanocarriers, vesicles, and emulsions [2,3,4,5,6] require multiple excipients and processing steps that increase cost. Furthermore, none suggest heat-stable oral insulin in a product format that is easily manufactured, is more accessible due to less stringent storage requirements, or has broader acceptability for all patient populations.

This study investigated the feasibility of a temperature-stable, gastric-resistant formulation for sustained oral delivery of insulin. We leveraged published data [7] and hypothesized that coupling insulin with bile salt as a lipophilic ion-pair would enhance intestinal transport, where encapsulation of this ion-pair within HP-β-CD will stabilize insulin, protecting from temperature and GI degradation. Furthermore, the cryoprotectant and polymer properties of HP-β-CD will allow insulin to be directly freeze-dried into a heat-stable tablet product.

## Research and methods

### Formulation

We evaluated the complexation efficiency of insulin with bile salt (sodium deoxycholate) and selected 1:28 as the optimal molar ratio for the formation of lipophilic ion-pair for complexation >95% insulin. The complex, which formed an insoluble precipitate, was mixed with a 2 wt% HP-β-CD solution and stirred for 30m at room temperature to form a stable colloidal solution.

### Tablet production

The colloidal solution was freeze-dried in blisters using established methods [8] to form tablets, followed by sealing the blisters (Figure 1).

**Figure 1.**
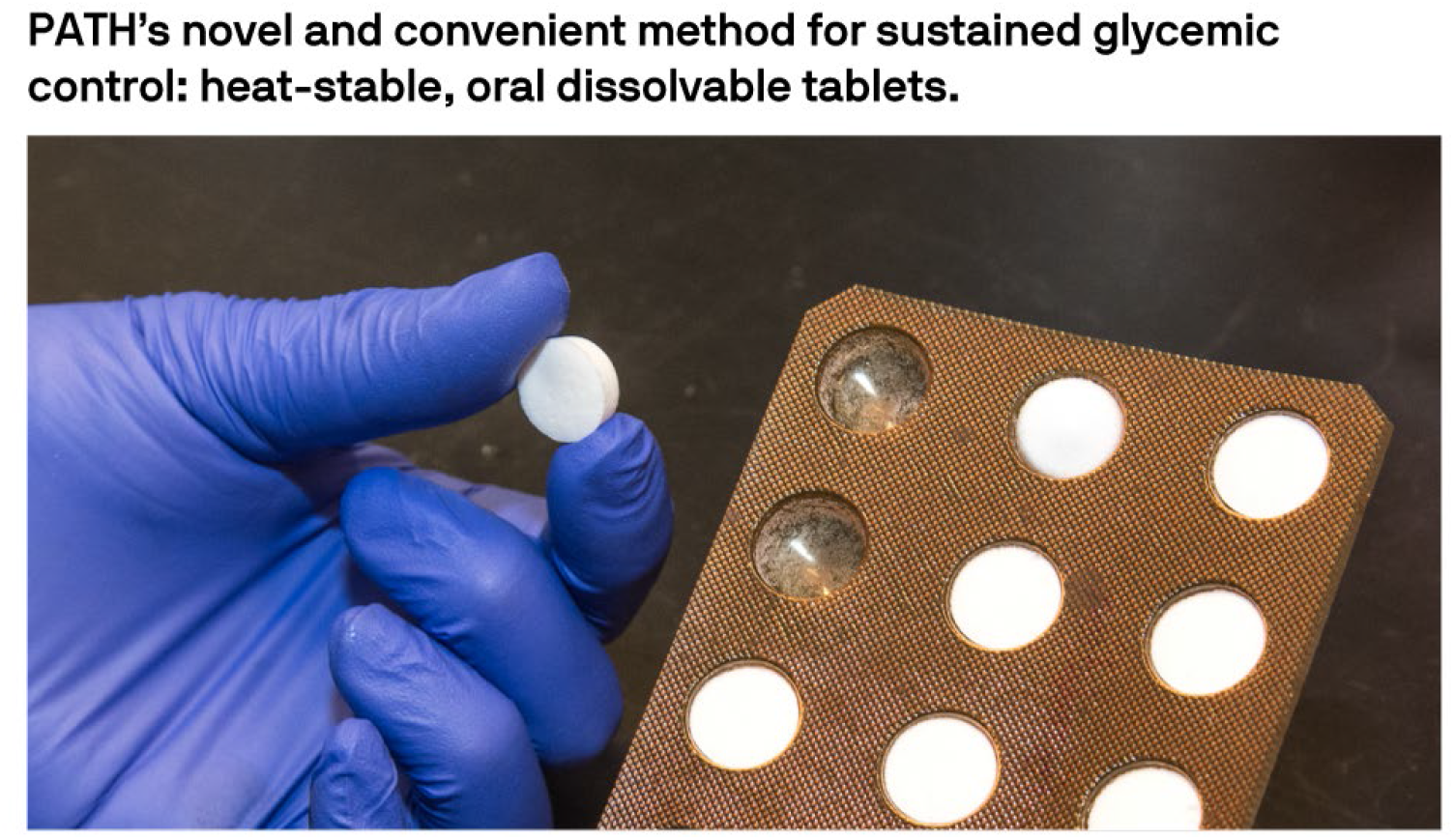
Novel freeze-dried, dissolvable tablet for oral and sustained delivery of insulin in convenient packaging for storage without cold chain.

### Three-month stability

Freeze-dried tablets and unformulated insulin control were stored at 2°–8°C, 30°C/65%RH, and 40°C/75%RH and tested for insulin content at week 0, week 2, week 4, week 8, and week 12 following the US Pharmacopeia specifications [9] for insulin to establish a sensitive analytical HPLC method for detection and quantification of insulin.

### In vivo evaluation

Eighteen (10 males, 8 females) Sprague Dawley rats with streptozotocin-induced diabetes, with >300 g/dl blood glucose received either subcutaneous insulin (control) or oral insulin tablet freeze-dried as an ion-pair encapsulated within HP-β-CD. Because the human oral insulin tablets were too large for oral administration to rats, crushed insulin tablets were delivered in gelatin capsules. Blood glucose and insulin levels in serum were tested at 0 min (pre-dose), 1h, 2h, 4h, 8h, 12h, and 24h post-dose, and once-daily body weights were measured. After the last timepoint, animals were humanely euthanized, without necropsy. In conducting research using animals, the investigators adhered to the laws of the United States and regulations of the Department of Agriculture.

## Results

Lipophilic insulin encapsulated in HP-β-CD retained 40% insulin content in simulated gastric fluid (SGF-USP) containing proteolytic enzymes [10], while liquid (unformulated) insulin was degraded and undetectable by the HPLC method. This formulation also maintained insulin content (90–110%) at 30°C/65%RH for 12 weeks and at 40°C/75%RH for 8 weeks by HPLC assay (Table 2).

As shown in Table 1, the test group receiving oral insulin (∼130 IU) showed a nearly 60% drop in glucose levels at 1h (500 mg/dL to 300 mg/dL) that was maintained even at 12h in rodents with diabetes, as compared to the subcutaneous control group. Glucose levels rose to ∼500 mg/dL by 8h in the control group, whereas a reduced glucose level was sustained at ∼300 mg/dL for both oral test groups. At 24h, the oral insulin tablet test group continued to maintain lower glucose levels than the subcutaneous control (data not shown).

**Table 1.** Stability of insulin content of thermostable oral insulin tablet and maintenance of lower glucose blood levels after administration in rats.

|  | Insulin stability: 30C, 65% RH |  |  |  |  |
| --- | --- | --- | --- | --- | --- |
|  | % insulin content |  |  |  |  |
|  | Time 0 | 2 weeks | 4 weeks | 8 weeks | 12 weeks |
| Insulin liquid control | 100 ± 0.01 | 85.1 ± 0.01 | 62.4 ± 0.3 | 41.2 ± 0.3 | 25.5 ± 0.7 |
| Oral insulin tablet | 97.8 ± 0.01 | 97.8 ± 1.9 | 95.1 ± 0.5 | 93.6 ± 1.7 | 86.9 ± 1.2 |
|  | Insulin stability: 40C, 75% RH |  |  |  |  |
|  | % insulin content |  |  |  |  |
|  | Time 0 | 2 weeks | 4 weeks | 8 weeks | 12 weeks |
| Insulin liquid control | 100 ± 0.01 | 49.5 ± 0.01 | 19 ± 0.1 | 3.7 ± 0.1 | 1.6 ± 1.6 |
| Oral insulin tablet | 100 ± 0.01 | 94 ± 0.2 | 91.2 ± 0.6 | 86.8 ± 0.8 | 79.1 ± 1.3 |
|  | Glucose blood levels (mg/dL) in rats after administration of insulin |  |  |  |  |
|  | Hours post dosing |  |  |  |  |
|  | Time 0 | 1 hour | 2 hours | 4 hours | 8 hours |
| SQ insulin control (13 IU) | 550 ± | 131.8 ± 59.6 | 185.8 ± 95.3 | 453.2 ± 164.6 | 471.0 ± 122.8 |
| Oral insulin tablet (130 IU) | 502.3 ± 94.8 | 283.8 ± 172.2 | 389.3 ± 242.4 | 373.7 ± 212.9 | 340.5 ± 164.7 |

SQ: subcutaneous injection; IU: international units.

## Discussion

The oral bioavailability of insulin is reportedly low primarily due to a) acid and enzymatic degradation encountered in the stomach and b) poor intestinal absorption. In our novel formulation, we formed a lipophilic ion-pair of insulin using bile-salt to facilitate insulin transport into the blood, followed by use of the inclusion properties [11,12] and cryoprotectant [13,14] attributes of HP-β-CD polymer in encapsulating and stabilizing the lipophilic insulin ion-pair to form a heat-stable oral tablet using a freeze-drying process. Our simple and unique approach, containing only two excipients, uses the natural ability of HP-β-CD [15] to encapsulate the lipophilic insulin ion-pair and maintains insulin stability without additional excipients or enteric coating of tablets.

Our hypothesis is that bile acid, ion-paired with insulin will modulate intestinal tight junctions, where the bile transporting enzymes will potentially transfer insulin across the intestinal epithelium. We tested this hypothesis in ex-vivo tissue studies conducted using Mattek Ep-intestinal tissue by measuring changes in transepithelial electrical resistance (TEER) across the intestinal tissue upon treatment with insulin (control) versus insulin-ion-pair FDT. Results from the TEER and tissue permeability evaluation indicated that ion-pair complexation of deoxycholate bile salt with insulin showed a greater drop in tissue resistance, likely due to the amphiphilic nature of bile salt, which resulted in making the tissue layer more fluid, thus decreasing the resistance. This finding also correlated well with the tissue permeability data where higher permeability of insulin (>50%) was observed across the tissues treated with ion-pair FDTs versus unformulated insulin control (data not shown).

In diabetic rodents, within 1h, the glucose levels were lowered by ∼60% and maintained these lowered levels for 12h; the slow, sustained level of glucose suggests encapsulation of insulin within the inclusion polymer HP-β-CD, which played a key role in the GI protection and release kinetics of insulin in this study. Additional optimization of the encapsulating polymer HP-β-CD in the formulation could reduce degradation and further lower the oral insulin dose required for glycemic control. This effect of encapsulation is also validated in the stability study: the encapsulated insulin was protected from temperature-induced degradation, while unformulated insulin (control) lost >50% content within 2 weeks under similar storage conditions.

These results suggest that oral, heat-stable insulin has the potential for further development into a sustained-release product capable of providing a steady, low level of insulin for maintaining blood glucose levels. Our tablets demonstrated robustness and superior thermostability compared to liquid insulin control, and retained potency without refrigeration, making it suitable for low- and middle-income countries where prevalence of diabetes is high and rising [16] and where reliable cold chain may not be available. Additionally, the ease-of-use of our freeze-dried oral insulin tablet has the potential to improve the management of diabetes for special populations including children and people with swallowing difficulties. Furthermore, the heat-stable tablets are produced using a well-established pharmaceutical manufacturing platform, enabling a rapid development timeline and potential to reduce insulin distribution costs.

The authors recognize the limitations of this study and seek to further evaluate the effect of HP-β-CD encapsulation in improving the absorption of oral insulin at lower doses. Additional work is warranted in several areas—including the degree of protection and bioavailability of insulin of the HP-β-CD inclusion polymer using a type 1 diabetes transgenic mouse model for pharmacokinetic/pharmacodynamic studies—to validate and advance this approach for oral, sustained delivery of insulin. Although the findings still need to be validated, we feel that it is important for the researchers working in this field to be aware of this innovative, yet simple formulation approach for oral delivery of insulin which if successful, has the potential to ease the diabetes burden.

## ETHICAL STATEMENT

### Ethics approval and consent to participate

In conducting research using animals, the investigators adhered to the laws of the United States and regulations of the Department of Agriculture. All animal research was conducted under Noble Life Sciences IACUC approval (NLS-570) dated 20-December-2019.

### Consent for publication

All authors have approved the manuscript and provide consent to its publication.

### Availability of data and materials

Data and materials are available by the authors by request.

### Authors’ contributions

All authors contributed to the manuscript composition and revision and approved the final draft.

## Acknowledgements

We thank Marge Murray and Abra Greene for assistance in preparing this manuscript.

